# Behaviour, biology, and evolution of vocal learning in bats

**DOI:** 10.1101/646703

**Authors:** Sonja C. Vernes, Gerald S. Wilkinson

## Abstract

The comparative approach can provide insight into the evolution of human speech, language, and social communication by studying relevant traits in animal systems. Bats are emerging as a model system with great potential to shed light on these processes given their learned vocalisations, close social interactions, and mammalian brains and physiology. A recent framework outlined the multiple levels of investigation needed to understand vocal learning across a broad range of non-human species including cetaceans, pinnipeds, elephants, birds and bats. Herein we apply this framework to the current state of the art in bat research. This encompasses our understanding of the abilities bats have displayed for vocal learning, what is known about the timing and social structure needed for such learning, and current knowledge about the prevalence of the trait across the order. It also addresses the biology (vocal tract morphology, neurobiology, and genetics) and phylogenetics of this trait. We conclude by highlighting some key questions that should be answered to advance our understanding of the biological encoding and evolution of speech and spoken communication.

## Introduction

Vocal production learning (herein ‘vocal learning’) is widely studied given its potential to shed light on the evolution of sensorimotor pathways, vocal-motor control, auditory-motor integration, and human speech. Without the ability to learn to modify our vocal output, humans would not have the extraordinary repertoire used to communicate via spoken language. A range of animals share this trait with humans, including some bird, bat, elephant, cetacean and pinniped species [1-3]. Utilizing multiple independent lineages in comparative studies has the potential to reveal critical steps necessary for biological encoding and evolution of this trait [4-6].

Bats provide a number of advantages compared to other vocal learning species. While birds are some of the most accomplished and well-studied vocal learners, Aves is separated from Mammalia by ∼300 million years of evolution, leading to substantial morphological (syrinx vs. larynx), neurobiological (nuclear pallium vs. layered neocortex), and genetic differences. Unlike many terrestrial mammals, bats are highly vocal and often have extensive vocal repertoires [7-11]. Many are also highly social [12] and exhibit various degrees of social complexity [13]. Other mammalian vocal learners are not as easily used for experimental studies, given their size and habitats. This is particularly true given that some bat species can be kept in breeding colonies in which they will reproduce. Group-living species present opportunities for controlled behavioural, morphological, neurobiological, and genetic studies to be performed with reasonably large numbers. Moreover, many bat species can live for a decade or longer in captivity [14], which permits longitudinal studies.

It should be noted, however, that studying bats also presents challenges. Despite tremendous diversity, not all species are easily bred in captivity (exceptions include frugivores, such as *Rousettus aegyptiacus, Phyllostomus discolor*, or *Carollia perspicillata,* or sanguivores, such as *Desmodus rotundus*). Furthermore, most bats have relatively long generation times (e.g. first reproduction occurs after either 1 or 2 years of age) and low fecundity (e.g. one or two offspring per year), meaning that the number of related animals per experiment will be necessarily low compared to mouse or some bird studies. Furthermore, because of the relatively small number of studies on bat vocal learning [4] many of the tools needed to investigate this trait are not yet present. Indeed, bat social communication calls have historically been relatively neglected, compared to work on echolocation, and thus the knowledge of and range of tools for assessing bat social calls is still limited. Similarly, comparatively little is known about detailed cortical structures of bats outside of auditory areas, which have been intensively studied, but only in a handful of echolocating species. Lastly, we must not overlook the importance of conservation concerns related to invasive bat research. Of the ∼1300 bat species identified, around 23% are considered near threatened, vulnerable, endangered or critically endangered, and a further ∼17% are data deficient (IUCN Red List; www.iucnredlist.org). Thus for ∼40% of bat species, we must approach their study with caution to avoid deleterious effects on populations.

Herein we summarise the state-of-the art of bat vocal learning, addressing the ‘WHAT’, ‘WHEN’, ‘HOW’, ‘WHO’ and ‘WHY’ of vocal learning in bats (Figure 1) [4], and highlight areas most in need of future study.

**Figure 1.**
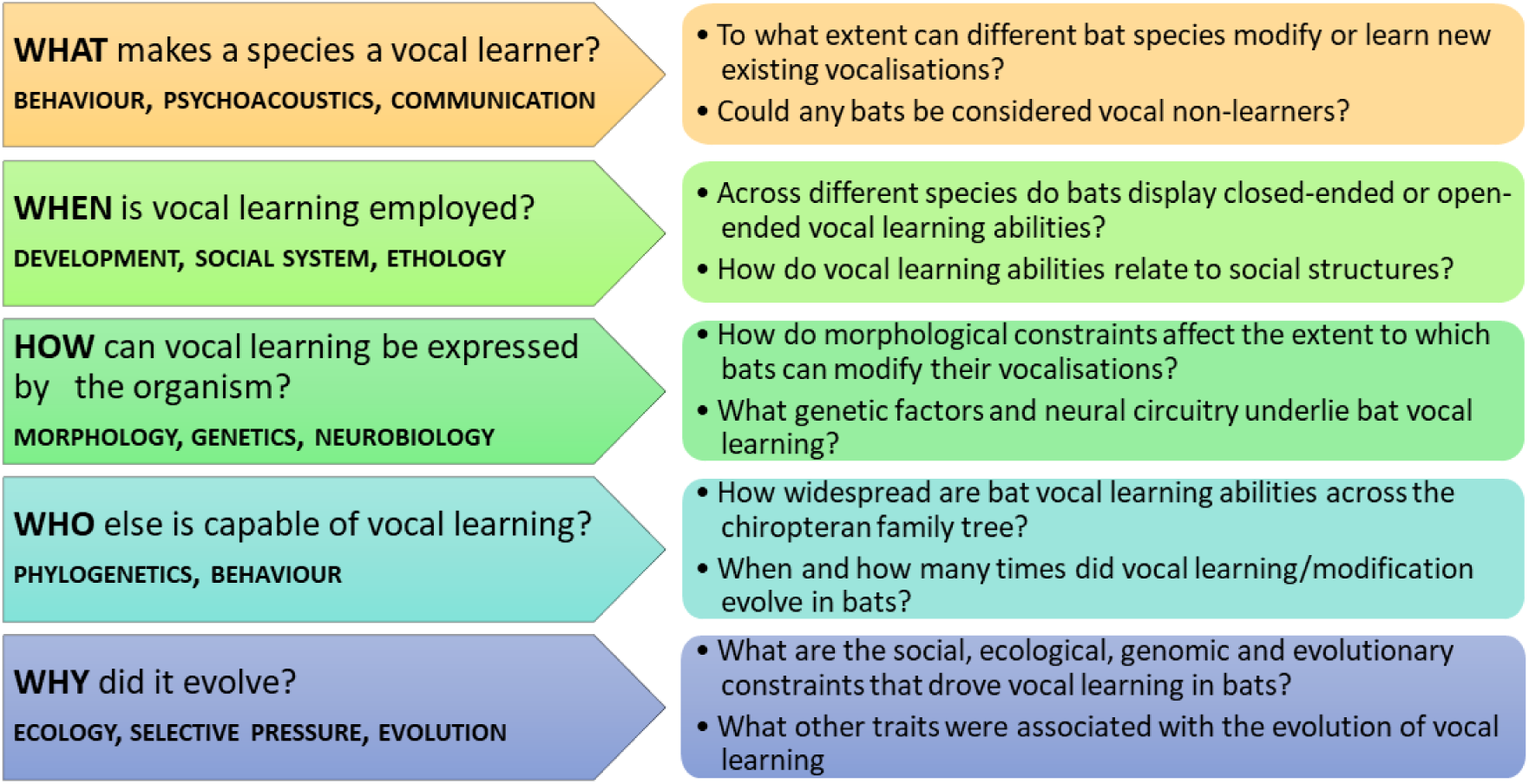
Framework for studying vocal learning. [4]. Left panels identify overarching questions at each level and the associated fields of study. Right panels indicate some key questions within each field that are a priority to address in bats.

### Evidence of vocal learning in bats (WHAT and WHEN)

We focus on vocal production learning [15] because of its importance for human spoken language, and because most studies on bats that claim to have relevance for vocal learning involve modification to vocalizations. In addition to reviewing the evidence for vocal learning, we discuss information regarding the ontogeny of vocal learning and the extent to which learning is or is not open-ended.

Definitions of vocal production learning vary from modification of a vocalization in response to auditory experience [15, 16], to development of calls that match those of individuals with whom the learner has interacted [17], and may be copied from conspecifics (imitation) or heterospecifics (mimicry). A proposed categorisation involves separating animals into limited versus complex vocal learners [17]. Limited vocal learning is described by Tyack (2016) as modification of acoustic features of existing vocalisations, while complex vocal learning refers to cases where an animal learns to produce a novel call type rather than modification of a pre-existing call. Another classification system is the continuum hypothesis, which proposes that vocal learning abilities occur along a continuum of abilities from limited (subtle modification of existing calls) to extensive (imitation of novel sounds) [18]. In both systems, birds, such as mynahs, parrots, or lyrebirds, that imitate the sounds of other species are clear examples of complex/extensive vocal learning, while species that learn to produce variants of a species-typical song, such as white-crowned sparrows, are considered more limited vocal learners. Importantly, even in sophisticated vocal learners, some part of the vocal repertoire may be innate - for example humans retain innate vocalisations, such as crying and laughter [17].

To date, all studies that have reported vocal learning in bats involve the modification of vocalisations. These vocalisations vary between species and include echolocation calls as well as social calls used either for parent-offspring reunions, territorial defense, or maintaining group integration (Table 1; Figure 2). We focus on examples that involve changes in vocalisation frequency, because such change is essential for speech production and rare in nonhuman mammals [19].

**Table 1.**
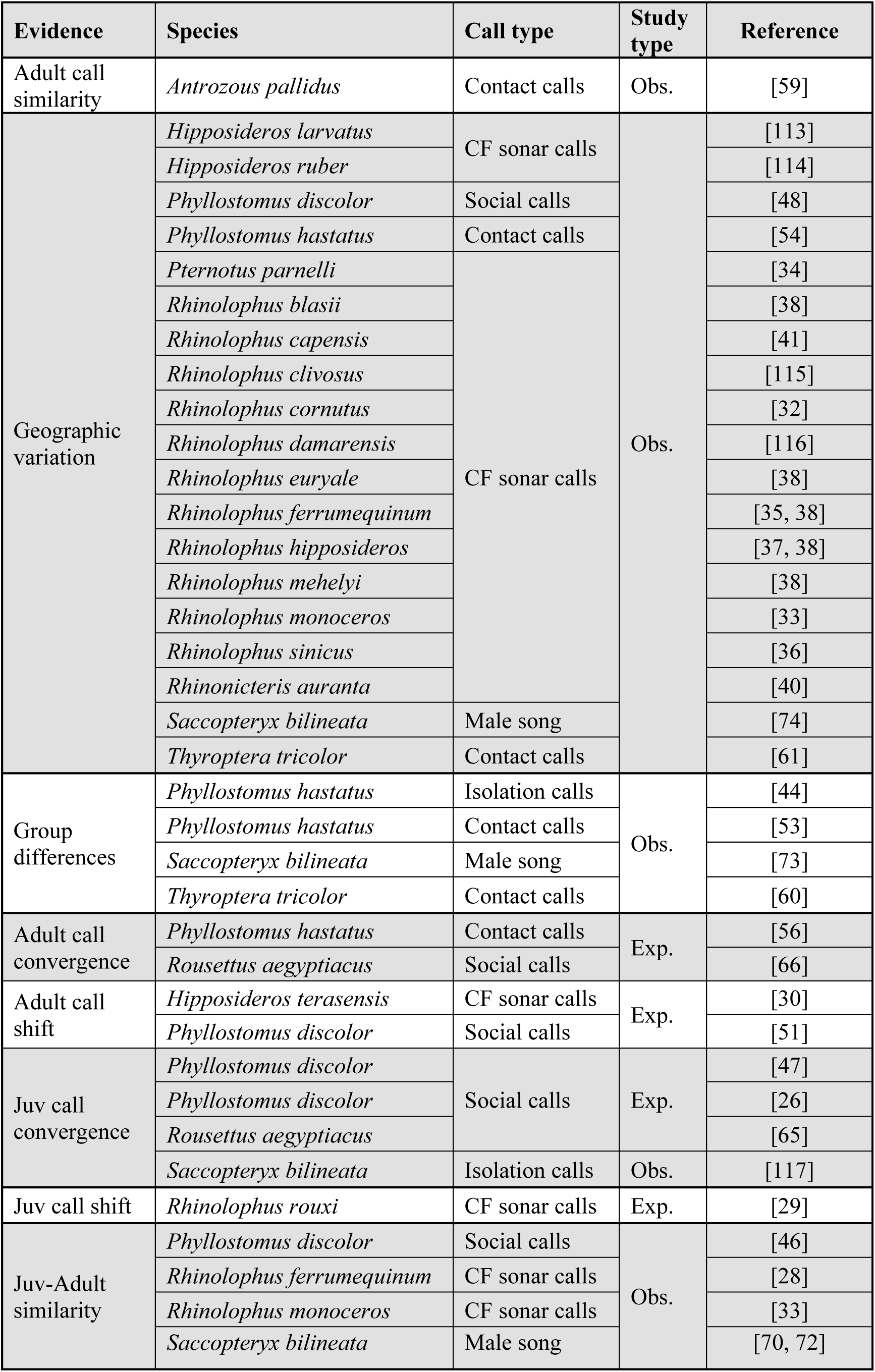
Evidence consistent with learned vocalisations in bats.

**Figure 2.**
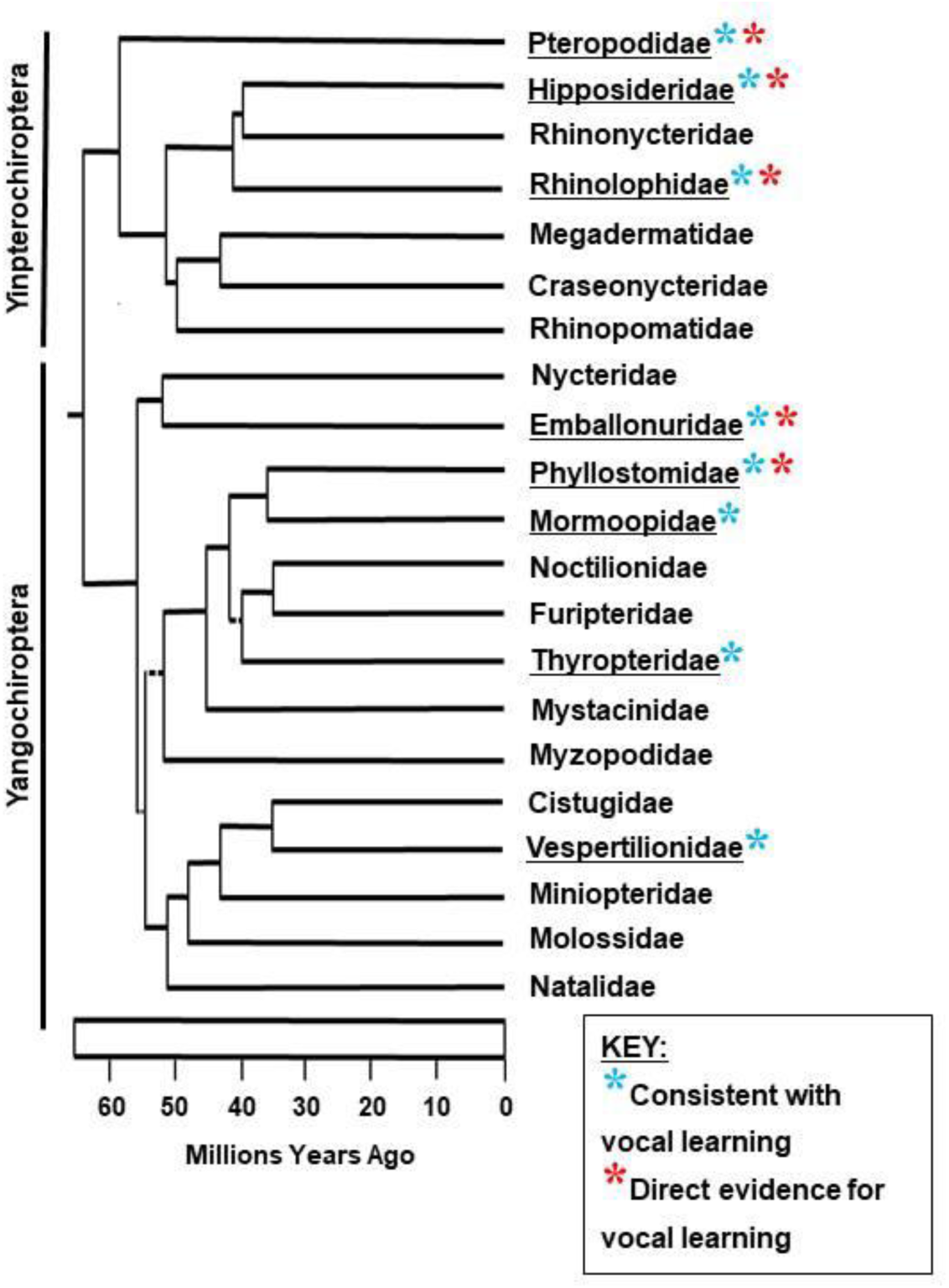
Evidence for bat vocal learning, mapped to the bat family tree. Behavioural studies relevant for vocal learning (see Table 1) have been categorised as ‘Consistent with vocal learning’ (found within eight families) or ‘Direct evidence for vocal learning’ (found within five of those families). ‘Consistent with vocal learning’ = adult call similarity, geographic variation, group differences, juvenile-adult similarity. ‘Direct evidence for vocal learning’ = adult call convergence, adult call shift, juvenile call convergence, juvenile call shift. Tree adapted from [106].

Detecting change in the frequency of echolocation calls can be difficult when bats emit calls that vary in frequency depending on the type of background clutter encountered. Consequently, there are few examples of bats that emit frequency-modulated (FM) echolocation calls to match conspecific calls. However, some FM bats clearly do respond to auditory feedback from conspecifics either by not producing echolocation calls [20] or by modifying call frequency in the presence of conspecifics [21-24] or ambient noise [25, 26]. In contrast, bats that rely on constant frequency (CF) echolocation calls produce long duration sounds that are centered on a narrow frequency band, so it is relatively easy to determine if the frequency of an echolocation call emitted while the bat is at rest is the same or different than another individual. Bats that use CF echolocation calls can rapidly modify their outgoing frequency so that the echo from a moving prey item consistently returns at the frequency of best hearing [27]. While such Doppler-shift compensation (DSC) clearly represents vocal modification in response to auditory feedback, this type of change is generally not considered vocal learning, because it is very rapid and transient, and evidence implicating learning is lacking. However, it is worth noting that bats that use DSC can clearly alter vocalisation frequency and might, therefore, be good candidates for vocal learning.

Indeed, one of the first examples of bat vocal learning involves change in the resting frequency (RF) of the constant frequency (CF) portion of the echolocation call of greater horseshoe bats, *Rhinolophus ferrumequinum*. Jones and Ransome [28] discovered that the RF of female bats change with age and that pups emit calls that match the RF of their mothers, suggesting that RF is at least partially the result of maternal transmission. This suggestion is consistent with an earlier study, which found that horseshoe bats deafened at 5 weeks of age eventually produced calls with RF up to 14 kHz different than normal [29]. A more recent study on captive Taiwanese leaf-nosed bats, *Hipposideros terasensis*, found that RF is influenced by the presence of conspecifics in the roosting group. Bats that moved into a colony adjusted their RF by as much as 4 kHz in 8 to 16 days to match the colony [30]. This study provides evidence that conspecific frequency matching is not limited to juveniles and indicates that the RF of bats in a colony could be influenced by conspecific matching, although stress induced changes to RF should be ruled out in the future.

The possibility that geographically isolated colonies of CF bats might then differ in RF by chance, similar to how differences in killer whale dialects have been hypothesized to arise [31], has been described as “cultural drift” and used to explain geographic differences in the RF of many CF bats [e.g. 32, 33-36]. Geographic variation in the RF of CF bats is widespread [37-39] and in some cases has been associated with morphological changes in nasal cavities [40, 41] indicating that some frequency differences may have a morphological and presumably genetic basis. In the absence of experimental data, it is difficult to distinguish genetic from learned transmission; thus, geographic variation in call frequencies provides, at best, indirect evidence for vocal learning.

A second example of vocal learning involves social calls used for parent-offspring reunions. Virtually all mammalian offspring emit calls when separated from their mother. In bats, these “isolation” calls typically have individually distinctive features [42], which are heritable [43, 44], and facilitate recognition and reunions [45]. In some species, mothers respond to isolation calls by emitting a “directive” call. In the lesser spear-nosed bat, *Phyllostomus discolor,* sinusoidal frequency-modulated isolation calls converge on a maternal directive call [46] or a simulated call if pups are hand-reared [47]. Maternal directive calls from different populations also contain group-specific frequency differences [48]. Using an operant conditioning paradigm developed initially for parrots [49, 50], Lattenkamp et al. [51] elicited spontaneous social vocalisations from isolated adult males, most of which resembled sinusoidal directive calls. By rewarding bats that met a highpass criterion, they were able to shift the duration of the bats’ calls by 20% and the frequency by up to 3% (0.45 kHz). Thus, *P. discolor* can modify the frequency of at least one type of social call both as juveniles and as adults.

The greater spear-nosed bat, *Phyllostomus hastatus,* also exhibits vocal learning, but for a different type of call. Adult females use loud, low frequency broad-band calls to coordinate foraging among group members [52]. These calls do not contain individually distinct features, but they do differ between groups within and between caves [53, 54]. Females disperse from their natal groups and then join groups containing unrelated individuals [55]. Adult vocalisations that contain group-specific information therefore require call modification based on experience with the conspecific group. Experimental transfer of individuals indicated that calls of all group members modified after transfer and converged within six months [56].

Indirect evidence for vocal learning of group integration calls has been reported for two other species. Female pallid bats, *Antrozous pallidus,* emit calls while flying prior to selecting a diurnal roosting site. These calls resemble isolation calls [57] and have individually distinct acoustic features [58]. In addition, call similarity among females is influenced both by relatedness and matriline, with the latter effect consistent with vocal convergence due to natal philopatry [59]. Similarly, disc-winged bats, *Thyroptera tricolor,* emit two types of calls much like a “Marco-Polo” game to locate group members in a furled leaf roost. One of the calls contains information about individual and group identity [60] and also exhibits geographic differences [61]. Both of these examples are consistent with vertical transmission by learning, but experiments are needed to rule out genetic transmission.

Another example of vocal learning involves Egyptian fruit bats, *Rousettus aegyptiacus*, a species in the family Pteropodidae, which uses lingual echolocation (tongue clicks) to enter and roost in caves and while foraging [62, 63]. This species has multiple call types in its repertoire [10] and often emits vocalisations during agonistic interactions in the roost [64]. Bats reared alone with their mothers take 5 months longer for their calls to converge on the spectral norm of adults than bats reared in a group [65]. Moreover, pups raised in the presence of calls broadcast at either higher or lower frequencies than normal, shifted their call frequencies toward those of the playbacks as they developed from 3 to 6 months of age [66].

The final example of vocal learning involves vocalisations given by greater sac-winged bats, *Saccopyteryx bilineata*, at a roost site. Males of this species produce elaborate multi-syllabic songs to attract and defend groups of females [67-69]. Young bats have more variable songs than adults [70], similar to songbird subsongs. Females disperse, so males within a colony tend to be related [71]. Features of some calls within a song converge on those of the territorial male, independent of relatedness, which is indicative of call matching [72, 73]. Some call features also differ between colonies [74], which is consistent with vocal learning, but could also be a consequence of male philopatry and genetic variation.

It is tempting to expect vocal learning to occur in bats that produce courtship songs given their apparent similarity to song in songbirds. Indeed males produce complex courtship songs in several other bat species, especially in the family Molossidae [9]. While courtship songs of at least one species, *Tadarida brasiliensis*, appear to have a hierarchical structure composed of composite elements [75] that can vary depending on context [76], population differences indicative of dialects or other evidence suggesting that these songs are learned have not yet been found [75]. However we would be remiss if we failed to point out a key difference in parental care between songbirds and most bats. Male songbirds typically sing at or near a nest where chicks are fed by both parents. Consequently, young birds are frequently exposed to male song. In contrast, male bats often sing on territories [77] that are far from the roost sites where pups are nursed by females. Sac-winged bats are an exception that supports this idea, because males sing at the roost where females nurse their young. We predict that if other cases of song learning are found in bats, they may be in those species where male song is heard by young.

Based on current evidence, call matching appears to occur during pup development of several distantly related species, e.g. *Rhinolophus ferrumequinum, Phyllostomus discolor*, and *Rousettus aegyptiacus*. In each case, the frequency of the call may change by a few kHz, but the type of call is characteristic of the species. In a few additional species, such as *Phyllostomus hastatus, P. discolor,* and *Hipposideros terasensis*, spectral matching also occurs in adult animals. We know too little at present to speculate on how widespread this type of adult vocal flexibility may be. Furthermore, we are not aware of any studies in which novel sound imitation or mimicry has been tested in bats. From observational studies this does not appear to be a common or obvious feature of bat vocal learning, however we can currently only speculate on the extent to which bats may be capable of producing completely novel sounds or sequences until it is more formally tested across species.

### Proximate mechanisms underlying vocal learning in bats (HOW)

Although the proximate mechanisms of vocal learning in bats are still relatively unknown, some morphological, neurobiological, and genetic features related to vocal learning have been explored.

Bats can produce calls across a much larger range of frequencies than most other mammals, including frequencies as low as 1 kHz (e.g. some social calls) and as high as 200 kHz (e.g. echolocation calls) [78]. Because the larynx and vocal tract shape these vocalisations, their morphology and neural control determine the range of possible vocalisations that an animal could produce. The structure of the larynx has been investigated in a number of echolocating bats and is broadly similar to that of other mammals [79, 80]. The source-filter model [19] initially proposed for human speech, applies to bats and other mammals; the laryngeal source and the vocal tract filter are considered to be largely independent [79, 80]. There are, however, some notable differences observed in the bat larynx. For example, echolocating bats display superfast laryngeal muscles that facilitate rapid call rates [81] and cartilage in the bat larynx ossifies earlier than in other mammals, which has been suggested to allow them to withstand the mechanical stress of echolocation [79]. The vocal folds of bats are comparable to other mammals, except that they feature an extended thin membrane that extends the vocal fold upwards. This vocal membrane is also found in some non-human primates [79, 80] and is thought to enable high-frequency or ultrasonic call production. Vocal membranes may also contribute to non-linear features during vocal production, such as subharmonics, independent pitch production and noisy call structures [80, 82]. In some rhinolophid bats, subglottal chambers have been observed [79, 83]. Lastly, in humans, contraction of both the cricothyroid and vocalis branch of the thyroarytenoid muscle contribute to fundamental frequency, but the fundamental frequency of echolocation calls in bats is mainly controlled by contraction of the cricothyroid muscle [79, 82].

While some features of calls may be determined by morphology, i.e. the structural and biophysical properties of the larynx, neural control is crucial for the tuning and spectro-temporal variations of the calls produced. As for other mammals, brainstem control of the laryngeal muscle is mediated via motor neurons projecting from the nucleus ambiguous [84]. In the midbrain, stimulation of the periaqueductal grey (PAG) has been shown in many mammals, including bats, to elicit vocalisations [85]. In bats echolocation calls and communication calls can be elicited via stimulation of distinct regions of the PAG [86]. Stimulation of other midbrain structures, including the paralemniscal area (PLA) and deep layers of the superior colliculus elicit echolocation, but not communication, calls in bats [84, 87]. In the bat cortex, stimulation of the anterior parts of the anterior cingulate cortex (ACC) can elicit echolocation, while stimulation of posterior regions of the ACC can result in vocalisations akin to communication calls [84, 88]. Although the complete vocal-motor control circuitry is not yet defined in bats, these data suggest that there may be different (but overlapping) neural pathways underlying echolocation and social call production. This is perhaps not surprising given the different primary purposes (communication vs. orientation), behavioural contexts, demands on sensorimotor integration, and spectrotemporal adjustments needed for these different call types [84].

In humans, voluntary vocalisations (e.g. speech) are controlled by the laryngeal motor cortex (LMC), located in the ventral primary motor cortex [89, 90]. The human LMC sends direct projections to the motor neurons of the nucleus ambiguus (NA) that control the laryngeal muscles [91, 92], as well as indirect NA connections via the basal ganglia (which is thought to play a role in control of learned vocalisations) [90, 93]. The Kuypers-Jurgens hypothesis proposes that the direct connection between the LMC and laryngeal motor neurons is required for vocal learning [94]. The presence of this direct connection is supported in vocal learning species, such as humans and songbirds, while data from vocal non-learning species, such as mice and non-human primates, suggest this connection is not present, or only very weakly connected [89, 90, 95-97]. The location of the laryngeal motor cortex in bats and the presence or absence of a direct connection between LMC and NA are yet to be determined, although work is underway in multiple labs and species to do so.

Work in songbirds [98] has suggested there are deep homologies in the molecular mechanisms underlying vocal learning across songbirds and humans. The most prominent gene involved in human speech disorders is *FOXP2*^*1*^. Disruption of this gene leads to childhood apraxia of speech (CAS) in which the spoken language of affected children is severely impaired [100]. In songbirds, FoxP2 disruption targeted to area X of the song learning circuitry (analogous to the mammalian anterior striatum) prevents accurate song learning in juvenile birds [101]. Even in mice, a vocal non-learning species, Foxp2 has related functions as heterozygous loss of Foxp2 impairs motor-learning skills [102]. Together these data suggest that some aspects of FoxP2 function in motor/sensorimotor learning is shared across these diverse lineages.

The sequence of the FoxP2 protein is highly conserved across evolution. At the amino acid level FoxP2 is 99.5% conserved between human and mouse [103] and 98.8% conserved between human and zebra finch [98]. FoxP2 in bats remains constrained (similarly displaying between 94.1-98.2% conservation with humans), however there is evidence for accelerated evolution of FoxP2 in Chiroptera. A survey of 13 bat species and 23 non-bat mammals, [104] identified more non-synonymous amino acid changes in bats compared to non-bat mammals. Although there is some uncertainty regarding the functional relevance of these changes, what seems clear is that bats have a higher diversity within the FoxP2 protein sequence than many mammals. Li et al. [104] suggest that this may be related to the emergence and diversity of echolocation in bats since the changes correlate with type of echolocation (e.g. the largely constant frequency producing bats of Yangochiroptera, vs. largely low duty cycle producing bats of Yinpterochiroptera, vs non-laryngeal echolocating fruit bats of Yinpterochiroptera).However, they also raise the possibility that this may be related to the vocal learning capacity of bats, given the role of FoxP2 in sensorimotor learning and rapid orofacial coordination, both of which are crucial to vocal learning and social communication. Importantly regulatory elements controlling the quantity or spacio-temporal expression of FoxP2 were not surveyed and these sequences are acknowledged as being increasingly important for evolution of complex traits. A recent non-coding effect on FoxP2 came from a derived human variant in intron 8 of *FOXP2*, not shared with Neanderthals, which was shown to influence expression levels [105]. The high quality genomes that are now being generated for diverse bat species [106] will give further insight into the coding and non-coding changes in *FoxP2*, and allow identification of other genes that may have contributed to vocal learning abilities in bats.

### When did vocal learning arise in the family tree (WHO)

The common ancestor of the Chiroptera arose approximately 60 million years ago [107]. Bats have colonised all continents except Antarctica, exhibit a range of social systems, and subsist on a variety of food sources (bats may be omnivores, frugivores, carnivores, or sanguivores). Current taxonomy recognizes 21 families. Among these, evidence consistent with vocal learning has been found in species from eight families with direct experimental evidence available for species in five of those families (Figure 2). The current distribution of vocal learning suggests three possibilities for its evolution: that it could have arisen in an ancestor and then been lost several times, that it evolved more than once, or since there is no confirmed vocal non-learning bat, it may have arisen once and still be present in all bats. These possibilities cannot be resolved without better determination of the extent of vocal learning among species and identification of which bats, if any, are vocal non-learners is essential. To date, no bat has been shown to have a purely innate repertoire or lack the ability to learn or modify their calls. Rather, there are bats that have not displayed obvious signs of vocal learning and thus are speculated to be possible vocal non-learners. However, the absence of evidence is clearly not evidence of absence, making additional behavioural studies crucial for understanding the evolution of this trait.

### What ecological and evolutionary factors led to vocal learning in bats – (WHY)

Multiple hypotheses have been proposed to explain how selection might favor vocal learning including group advertisement by vocal dialects, information sharing, sexual selection, environmental adaptation, and individual recognition [108]. These hypotheses are not mutually exclusive and might also be important in some lineages but not others. However, we think that much of the current evidence, both indirect and direct, indicates that vocal learning in bats involves modifications to calls that are used for group integration and/or recognition. The functional significance of being able to recognize a group member likely varies among species, but in many cases, it could be very important for locating an appropriate roosting site [109, 110]. Opportunities for cooperation depend on associating with known individuals, which will be enhanced when individuals roost together over extended periods of time [13]. As noted above, while many male bats produce songs to attract or defend females much like songbirds [9], in most cases offspring do not develop near where males sing. Consequently, the sexual selection hypothesis seems unlikely to apply in most species.

To understand the social, ecological, and evolutionary forces that drove the appearance of vocal learning abilities in some lineages, but not others, a clear understanding of the typology of the trait, and its distribution across species is crucial. Although the current categorisations, such as limited vs. complex learning or the continuum hypothesis, are useful, they each have their limitations and do not fully encompass the complexity of phenotypes observed across vocal learning species. A precise classification system for vocal learning would allow us to identify specific biological features that accompany those abilities, and begin to address evolutionary questions. Although the need for these studies is not limited to bats, the possibility for more invasive studies in bats, for example tracing neural circuitry or manipulating gene activity, presents an opportunity to address the biology of vocal learning in a way not possible in most other vocal learning mammals, such as cetaceans or elephants. This would also allow comparisons with songbirds - the current dominant animal model of vocal learning - allowing us to ask if mechanisms identified in avian systems are convergent or divergent from those found in mammals. Comparing similarities and differences associated with vocal learning from birds, to bats, to humans will get us closer to understanding the evolution of this rare and complex trait.

### Concluding remarks and outstanding questions

The study of vocal learning in bats has been gaining attention in recent years. The resultant increase in researchers around the world studying the behavior, morphology, neurobiology, and genetics of bat vocal learning will lead to a better understanding of this trait, as well as generating better tools to allow us to address these questions in the future. Given the relatively early stage of bat vocal learning research (e.g. compared to songbirds) it seems timely to outline some key questions that the field needs to answer.

#### WHAT makes a species a vocal learner?

It is important that we determine the extent of vocalisation modification possible in bats, and whether any species can imitate or mimic novel vocalisations or sequences. Identification of bat species that are vocal non-learners would be very valuable. Ruling out vocal learning is a difficult task. To do this, it must be demonstrated that the vocal repertoire does not change as a consequence of auditory input or conspecific interaction. This could be determined by deafening, isolation, playback tutoring, or cross-fostering experiments; however, such experiments may not be feasible in some species.

#### WHEN is vocal learning employed?

Modification of bat vocalisations, as described herein, has been demonstrated in both juvenile and adult bats, suggesting some degree of open-ended learning. Further studies should determine the degree of open-ended learning across different species, and also if this relies on social interactions, such as specific adult tutors simply listening to conspecifics.

#### HOW is vocal learning expressed?

Technological advances are opening the way for in depth studies of the morphological, neurobiological, and genetic factors underlying bat vocal learning abilities. At the morphological level, biophysical studies of the larynx and vocal apparatus can give insight into social call production and the constraints of modification or imitation of calls. Tracing, electrophysiological, optogenetic, and neuroimaging techniques can shed light on the neural circuitry involved in vocal learning in the bat brain. Such studies can answer questions about the connectivity of a vocal learning brain, as well as whether bats have a direct connection between the cortex and laryngeal motor neurons. At a genetic level we can determine if there is a causative link between known language-related genes (such as FoxP2 [111]) by knocking down gene expression in the bat brain and observing the effect on behavior. Genomic and transcriptomic studies contrasting vocal learners and nonlearners will allow us to uncover new, potential causative genes as well as wider molecular networks that underlie bat vocal learning abilities.

#### WHO is capable of vocal learning?

Some degree of vocal learning has been identified in 8 of the 21 families of bats, however this still represents a very small fraction of all extant species. As such we need a much better understanding of how widespread bat vocal learning abilities are across Chiroptera to be able to determine how many times they likely have evolved.

#### WHY did it evolve?

Further behavioural, neurobiological, genetic, and phylogenetic investigations will give us clues to the social, ecological, and evolutionary constraints that drove vocal learning in bats. This should involve a determination of whether related traits were associated with the evolution of vocal learning (e.g. turn taking, echolocation, cooperation, timing, rhythmicity/synchronisation) [4, 9, 112].

Although there is a long road ahead to answer these questions, the multitude of approaches being undertaken by different groups around the world to address such questions gives reason to be optimistic about the contribution of bat research to the underpinnings of vocal learning, how it may have evolved in mammals, and the human ability for speech learning.

## Footnotes

1 Following standard nomenclature [99] Kaestner, K.H., Knochel, W. & Martinez, D.E. 2000 Unified nomenclature for the winged helix/forkhead transcription factors. *Genes Dev.* **14**, 142-146., human gene symbols are italicized and in upper-case (*FOXP2*), rodent gene symbols are italicized with only the first letter in upper-case (*Foxp2*) and other species, or a mix of species in upper and lower (*FoxP2*). Protein names are not italicized (FOXP2/Foxp2/FoxP2).

